# Nucleolus transformation and Pol1 activity pattern dual transcriptomes of neuronal subtype identity

**DOI:** 10.64898/2026.05.09.724016

**Authors:** Dmitrii Zagrebin, Stefan R. Stefanov, Nadine Brombacher, René Buschow, Lukas Behrens, Ekaterina Borisova, İbrahim A. Ilık, Beatrix Fauler, Thorsten Mielke, Tuğçe Aktaş, Mateusz C. Ambrozkiewicz, Helene Kretzmer, Andreas Mayer, Elke Deuerling, Matthew L. Kraushar

## Abstract

Gene expression directs neuronal fate during development with multiple RNA polymerases in the nucleus. mRNA transcribed by Pol2 ultimately depends on the rRNA transcribed by Pol1 in the nucleolus, which generates a ribosome pool to control mRNA translation. How Pol1 activity is synchronized with the Pol2 transcriptome to direct gene expression and neuronal fate is unknown. By creating a high-resolution map of nucleoli in the developing neocortex, we discover that the nucleolus is restructured in neuronal lineages with distinct Pol1 activity. With a new method for rRNA and mRNA dual-transcriptome sequencing in single-cells, we reveal that rRNA transcription is neuronal subtype and stage specific. Rather than a general feature of neural progenitor differentiation, rRNA production peaks transiently in post-mitotic subcortical projection neurons, which have atypical nucleoli containing a single, large cavity. Pol1 inactivation redirects these neurons towards a distinct fate that expresses a hybrid of subcortical and intracortical projection neuron markers. Thus, nucleolar architecture and Pol1 activity pattern dual rRNA-mRNA transcriptome signatures of neuronal subtype identity.

## Introduction

The neocortex is a layered brain region in mammals formed by the sequential generation of neurons in development ^1^. Neuronal subtypes are positioned in different neocortex layers, and project their axons to distinct targets in the nervous system ^2,3^. Timed gene expression specifies the identity of neuronal subtypes. Patterns of mRNA transcription and translation activity vary in neuronal lineages to generate neuronal subtype diversity ^1–7^. How mRNA transcription and translation are coordinated, and where their regulation intersects, are largely unknown in neocortex development.

The nucleolus is an organelle where transcription and translation regulation converge ^8–10^. As RNA polymerase II (Pol2) transcribes mRNA, RNA polymerase I (Pol1) transcribes rRNA in the nucleolus to generate ribosomes responding to mRNA translation regulation. Pol1 activity initiates ribosome biogenesis, which recruits hundreds of proteins to the nucleolus ^11^. Inside the nucleolus, ribosomal proteins bind stably to the rRNA scaffold forming the ribosome core, while assembly factors like Npm1 interact dynamically with emerging rRNA ^12–17^. The Pol2 transcriptome includes messages for ribosomal proteins that initiate future rounds of Pol1-driven ribosome assembly and translation regulation. Thus, gene expression regulation in neurodevelopment must engage cycles of mRNA and rRNA transcription, translation, and ribosome assembly in the nucleolus ^1^. However, how nucleolar Pol1 transcription of rRNA is synchronized with the expression of Pol2 transcriptomes in neuronal subtypes is unknown.

Here we discover dynamic nucleolar architectures that were previously not seen in neocortex neuronal lineages with a combination of light, electron, and expansion microscopy. To track the simultaneous activity of Pol1 and Pol2 transcription in neuronal lineages, we developed a single-cell RNA-seq method that captures full-length, high-fidelity rRNA and mRNA transcriptomes called RiboSTS (<u>Ribo</u>some <u>S</u>ignatures <u>T</u>argeted by high-throughput single-cell <u>S</u>equencing). RiboSTS analysis of neocortex development revealed that rRNA transcription is not a housekeeping function common to neuronal lineages. Instead, rRNA production is heterogeneous in subtypes of progenitors and differentiating neurons. rRNA expression peaks transiently in a differentiating lower layer neuron subtype, in contrast to low expression in all subsequent upper layer lineages. Knockdown of Pol1 in lower layer neurons leads to the co-expression of lower and upper layer subtype markers in the same cells. Taken together, nucleolar architecture and Pol1 activity are defining features of neuronal lineage-specific transcriptional programs and neuronal fate in the neocortex.

## Results

We first visualized nucleoli and their activity throughout mouse neocortex neurogenesis by immunolabeling with Npm1 ^12–17^ (**Fig. 1a, Extended Data Fig. 1**). Intense nucleoli are apparent in ventricular zone (VZ) progenitors and cortical plate (CP) neurons from E12.5 to E14.5. Strikingly, at E15.5 nucleolar intensity decreases sharply in VZ progenitors and post-mitotic neurons forming lower (LL) and emerging upper layers (UL). Quantification of Npm1 signal revealed a decrease in nucleolar intensity and size, but increase in number – from one nucleolus on average in E12.5 cells to five in LL neurons at E17.5 (**Fig. 1b**).

**Figure 1.**
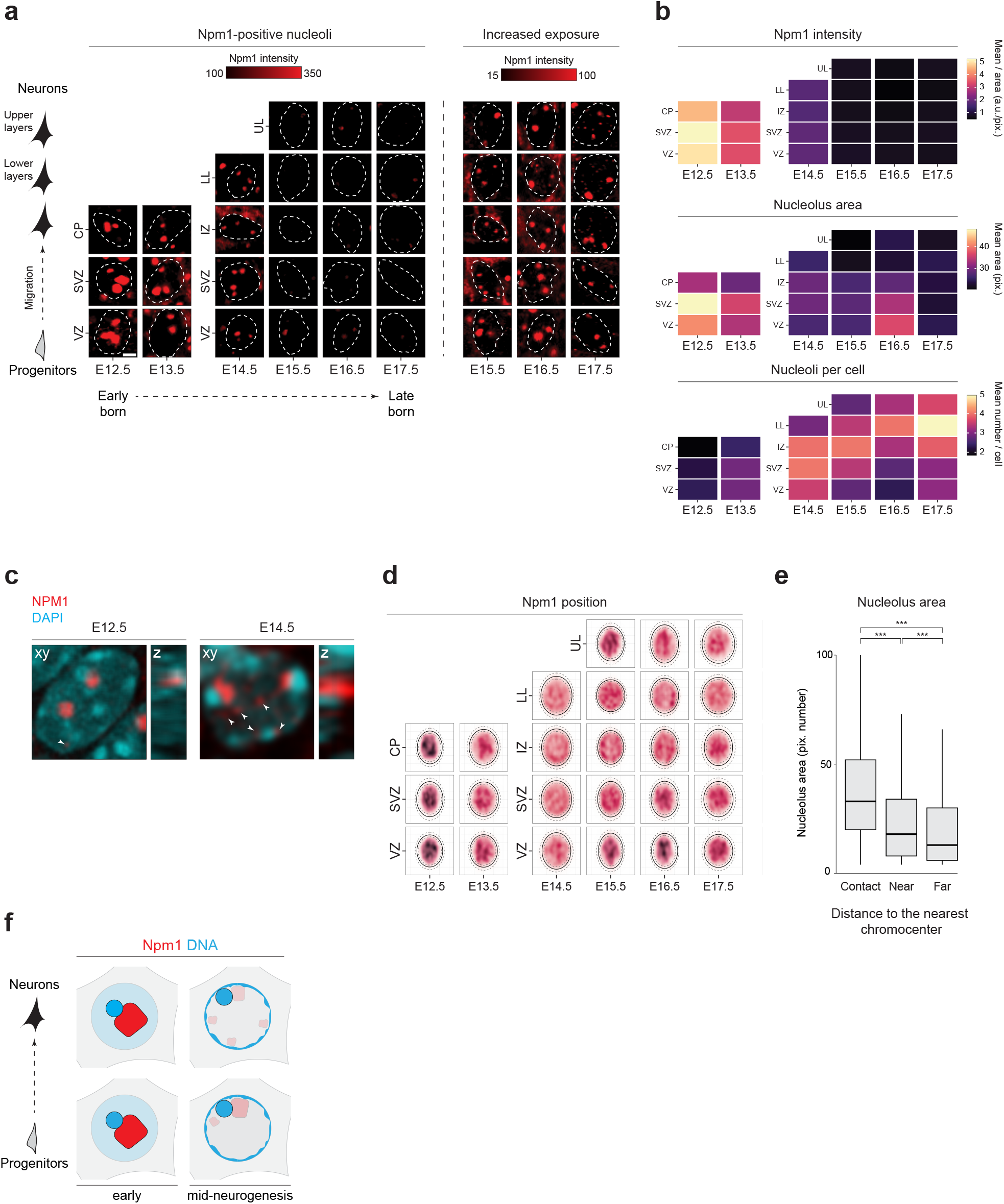
Nucleolar Npm1 depletion and mobilization to the nucleus periphery. **a**: Immunohistochemistry labeling of Npm1 (red) in coronal neocortex sections. Scale bar: 3 µm. Abbreviations: VZ: ventricular zone; SVZ: subventricular zone; CP: cortical plate; IZ: intermediate zone; LL: lower layers; UL: upper layers. **b**: Quantification of (a) for nucleolar Npm1 intensity, area, and number in neocortex cells. Multiple images taken of separate hemispheres from n = 3 biological replicates. On average, ~3000 cells were analyzed per heatmap tile. a.u.: arbitrary fluorescence units; pix.: pixel number. 1 pix. ≈ 0.22 µm. **c**: Representative images showing nucleolar fragmentation from stage E12.5 to E14.5. Chromocenters were defined as intense DAPI (cyan) puncta. White arrows indicate multiple small Npm1 foci. **d**: Quantification of (a) for Npm1 localization within the nucleus relative to the nuclear center of mass with an elliptical approximation of nuclear area. Solid line: median. Dashed lines: 25^th^ and 75^th^ percentiles. Kernel density estimates are shown (white, low density; dark red, high density), normalized within each panel (stage and layer). **e**: Nucleolus area categorized by distance to the nearest chromocenter. Distance categories – Contact (0–1 pix. or 0–0.22 µm), Near (1–5 pix. or 0.22–1.10 µm), and Far (>5 pix. or >1.10 µm). Statistical significance was assessed by one-way ANOVA followed by Dunnett’s multiple-comparisons test (vs. “Contact”). P values (***P < 0.001). **f**: Summary of lineage- and stage-dependent nucleolar dynamics and chromatin organization.

Nucleoli often contact “chromocenters” in mouse nuclei ^18^ (**Fig. 1c**). In mouse acrocentric chromosomes that contain nucleolar organizing regions, pericentromeric heterochromatin (chromocenters) are near rDNA arrays. Concurrent with a decrease in Npm1 intensity and increase in nucleoli, chromocenters and nucleoli move in tandem towards the nuclear periphery – especially at E14 in the intermediate zone (IZ) (**Fig. 1c-d, Extended Data Fig. 1-3**). DNA localization to peripheral lamina-associated domains in the nucleus typically reflects transcriptional repression ^19^. Chromocenters contacting nucleoli are particularly mobile, while those without nucleoli are consistently positioned at the periphery (**Extended Data Fig. 3b**). Nucleoli contacting a chromocenter are much larger than those without one (**Fig. 1e, Extended Data Fig. 3c**), suggesting that nucleolus-chromocenter structures may be more active sites of rRNA transcription ^20^. Thus, nucleoli are highly mobile, structurally heterogeneous, and are depleted for Npm1 in neocortex progenitors and neurons at mid-neurogenesis (**Fig. 1f**).

### Dynamic nucleolar architecture in neocortex neuronal lineages

The nucleolus is a dynamic organelle that changes its structure in a context-dependent manner, including transcription activity, stress, and development ^8,9^. The structure of the nucleolus in neocortex neurogenesis is unknown. We next mapped the interior structures of nucleoli with electron microscopy in the neocortex from E12.5-E17.5 (**Fig. 2a, Extended Data Fig. 4**). Major structural changes occur in the nucleolus interior throughout neurogenesis. Progenitors in the VZ harbor large nucleoli with many small phase-bright foci, typical of active nucleoli ^8^. However, as neuroblasts migrate into the subventricular zone (SVZ) and through the IZ, the nucleolus becomes dominated by a single, large, central cavity ~450nm wide. Cells with nucleolar cavities accumulate in lower layers of the cortical plate at E14.5, and persist until E16.5. At E17.5, when neurogenesis is largely complete and differentiation continues, nucleoli again transition towards a mixture of structures, suggesting a gradual reversion to prior nucleolar states.

**Figure 2.**
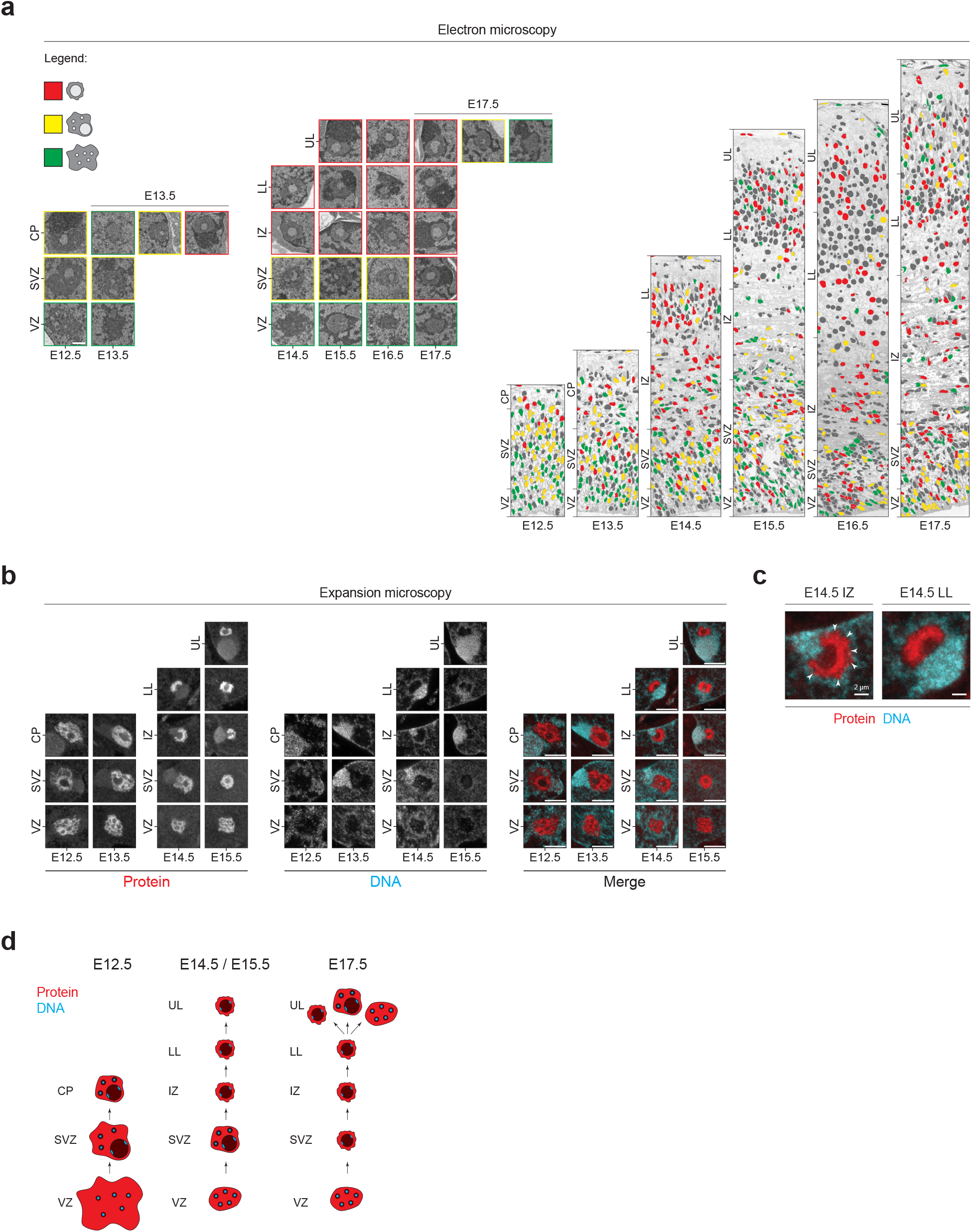
Map of the nucleolus interior and cavity formation in neuronal lineages. **a**: Electron microscopy in coronal neocortex sections. Left: Zoom-in of individual nucleoli. Colors categorize different nucleolar morphologies: canonical (green), intermediate (yellow), single nucleolar cavity (red). Scale bar: 0.75 µm. Abbreviations: VZ: ventricular zone; SVZ: subventricular zone; CP: cortical plate; IZ: intermediate zone; LL: lower layers; UL: upper layers. Right: nucleolus morphology categories mapped onto nuclei in complete coronal sections of the neocortex (gray: no nucleolus identified). **b**: Expansion microscopy in coronal neocortex sections with labeling of total protein (red) and DNA (cyan). Scale bar: 10 µm (scale post-expansion). **c**: Zoom of the E14.5 intermediate zone and lower layer nucleolus from (b). Arrows highlight collapsing protein structures. **d**: Summary of lineage- and stage-dependent nucleolar architecture.

To investigate the components within neocortex nucleoli, we performed expansion microscopy of the neocortex from E12.5-E15.5 with DNA and protein staining (**Fig 2b, Extended Data Fig. 4**). Expansion of neocortex tissue revealed complex nucleolar architectures, with dense protein and DNA staining that illuminate the structures seen by electron microscopy. Progenitors in the VZ contain honey comb-like protein density including puncta of DNA, characteristic of fibrillar centers where rDNA is transcribed ^8^. As neurons traverse the intermediate zone at E14.5, the honey-comb protein density begins to collapse (**Fig. 2c**). Most nucleoli in lower layer neurons at E14.5 contain a single rim of dense protein, with low protein and low DNA content in the cavity interior. The large cavities are reminiscent of “nucleolar vacuoles” in plants and *C. elegans* ^21–26^, “nucleolar bodies” or “No-bodies” in yeast ^9^, or possibly “A-bodies” ^27,28^. Such cavities open after intensive rDNA transcription or stress, are devoid of typical nucleolar proteins, and contain components for the surveillance of defective rRNA synthesis. To our knowledge, such nucleolar cavities are atypical of mammals, and not previously described in neurodevelopment. The Npm1 labeling and high-resolution imaging combined demonstrate that nucleoli remodel their architecture at mid-neurogenesis, which may reflect major changes in nucleolar Pol1 activity (**Fig. 2d**).

### A single-cell method to track rRNA transcriptomes throughout neurogenesis

We next aimed to track nucleolar Pol1 activity by measuring rRNA production in neocortex neuronal lineages. Methods to track rRNA production are lacking because rRNAs present several unique obstacles to single-cell sequencing (**Extended Data Fig. 5**). First, rRNAs are tightly bound to ribosomal proteins within large ribonucleoprotein complexes. Second, rRNA is highly structured, long, and GC-rich, which yields large gaps in coverage and prohibits accurate quantification. Third, rRNA lacks a 3’-poly-A tail commonly used to prime reverse transcription. Finally, rRNA constitutes the vast majority of total RNA, and is typically depleted during library preparation to permit sequencing of mRNA, which accounts for just 1-5% of the transcriptome. A technology that defines a cell by both its rRNA and mRNA transcriptomes would thus need to cope with a major imbalance in their stoichiometry. We systematically tackled these obstacles in the development of RiboSTS to capture the 18S rRNA and mRNA transcriptomes in single cells (**Fig. 3a, Extended Data Figs. 5-9**).

**Figure 3.**
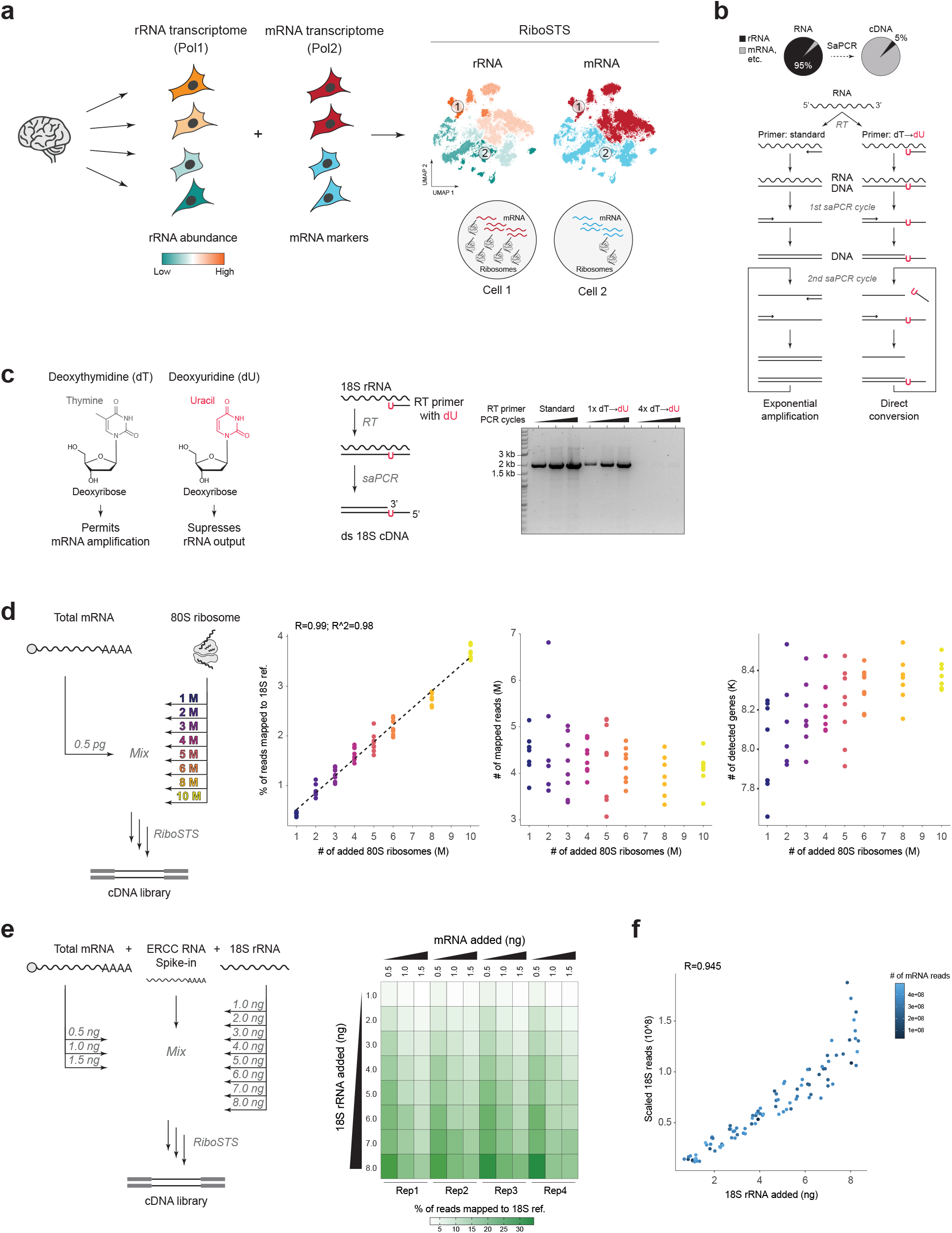
Dual rRNA-mRNA transcriptome sequencing in single cells with RiboSTS. **a**: Schematic of the single-cell features analyzed by RiboSTS. **b**: Flowchart illustrating the reaction to re-balance rRNA abundance via selective amplification PCR (saPCR). Deoxyuridine (dU) is utilized in the reverse transcription (RT) primer to prevent amplification of rRNA-derived cDNA. **c**: Left: comparison of deoxythymidine (dT) and deoxyuridine structures. Right: PCR products of reactions with varying dT-to-dU substitutions in the RT primer across increasing PCR cycles, analyzed by agarose gel electrophoresis. **d**: Left: RiboSTS quantification accuracy testing for the simulated physiologic range of rRNA/ribosomes in a single cell. A titration series of 1-10 million (M) ribosomes mixed with a constant 0.5 pg of mRNA isolated from HEK293 cells. Right: mapping statistics using a modified human genome reference. Shown are the percentage of fragments mapping to 18S rRNA correlated to the known ribosome input (Pearson R = 0.99), total RNA-seq fragment counts, and number of detected genes (complexity) per well. Points represent individual wells, colored by ribosome input. Note that <4% of total sequencing reads (y-axis) map to rRNA at the highest ribosome concentration. **e**: Left: RiboSTS quantification accuracy tested by varying the relative amounts of mouse mRNA and 18S rRNA against a constant ERCC RNA spike-in. Right: mapping statistics showing percentage of fragments mapping to 18S rRNA across technical replicates (Rep1-4). **f**: Mapping statistics of (e) showing linear correlation of ERCC-normalized reads mapping to 18S rRNA with added 18S rRNA amounts (Pearson R = 0.945). Points represent individual wells, colored by number of fragments mapping to mRNA.

A major innovation of RiboSTS is a method to balance the cDNA library when sequencing both rRNA and mRNA transcriptomes (**Fig. 3b, Extended Data Fig. 7**). Given that the physiologic range of ribosome number per cell is ~1-10 million ^29,30^, we reasoned that unlike mRNA, rRNA does not require amplification. Instead, a 1:1 conversion of 18S rRNA to cDNA generates a sufficient number of molecules for sequencing. To achieve this, we took advantage of the fact that family B DNA polymerases from the archaea, commonly used in PCR, strongly bind to uracil in the DNA template-strand and stall polymerization in response to this base ^31,32^. We hypothesized that substituting thymine with uracil (dT→dU) at the 3’ end of the 18S rRNA RT primer would not affect the reverse transcription (RT) reaction, but would halt DNA polymerase during PCR, preventing the synthesis of the RT primer binding site and thus circumventing 18S cDNA amplification. We call this method to selectively amplify the mRNA library while rebalancing rRNA library abundance “selective amplification” PCR (saPCR) (**Fig. 3b**). We performed RT on total RNA using 18S rRNA primers with varying numbers of dT-to-dU substitutions, followed by PCR amplification for different numbers of cycles (**Fig. 3c**). The uracil-induced stalling of DNA synthesis is cumulative, with at least four dT-to-dU substitutions fully suppressing 18S cDNA amplification (**Fig. 3c**), without impairing cDNA yield from mRNA (**Extended Data Fig. 7**).

To test whether the number of 18S cDNA sequencing reads correlates proportionally with the amount of rRNA in a cell, we simulated a single-cell reaction (**Fig. 3d**). The mRNA remained constant (0.5 pg) while varying the number of added 80S ribosomes (1×10^6^ – 1×10^7^), both within the physiological range for an average mammalian cell ^33,34^. The proportion of 18S rRNA reads showed a linear correlation with the number of ribosomes added. Importantly, rRNA reads constituted between 1% and 5% of total reads, and the number of ribosomes in the reaction did not compromise the complexity of sequencing libraries. Furthermore, RiboSTS is robust to changes in mRNA abundance simultaneous with changing rRNA abundance (**Fig. 3e-f**). Thus, RiboSTS captures the dynamic range of rRNA and mRNA transcriptomes by sequencing in single cells.

### Timed neuronal subtype-specific rRNA production in neocortex neurogenesis

We first employed RiboSTS to track rRNA production during the period of nucleolus restructuring in neocortex neuronal lineages, including ~4000 cells across four stages from E13.5-E16.5 (**Fig. 4a**). Cells clustered by embryonic stage, location, and subtypes of progenitors and post-mitotic neurons. These data are comprehensive for both apical (AP) and intermediate (IP) progenitors, which transition to subtypes of early-born lower layer and late-born upper layer subcortical and intracortical projection neurons (**Extended Data Fig. 10**).

**Figure 4.**
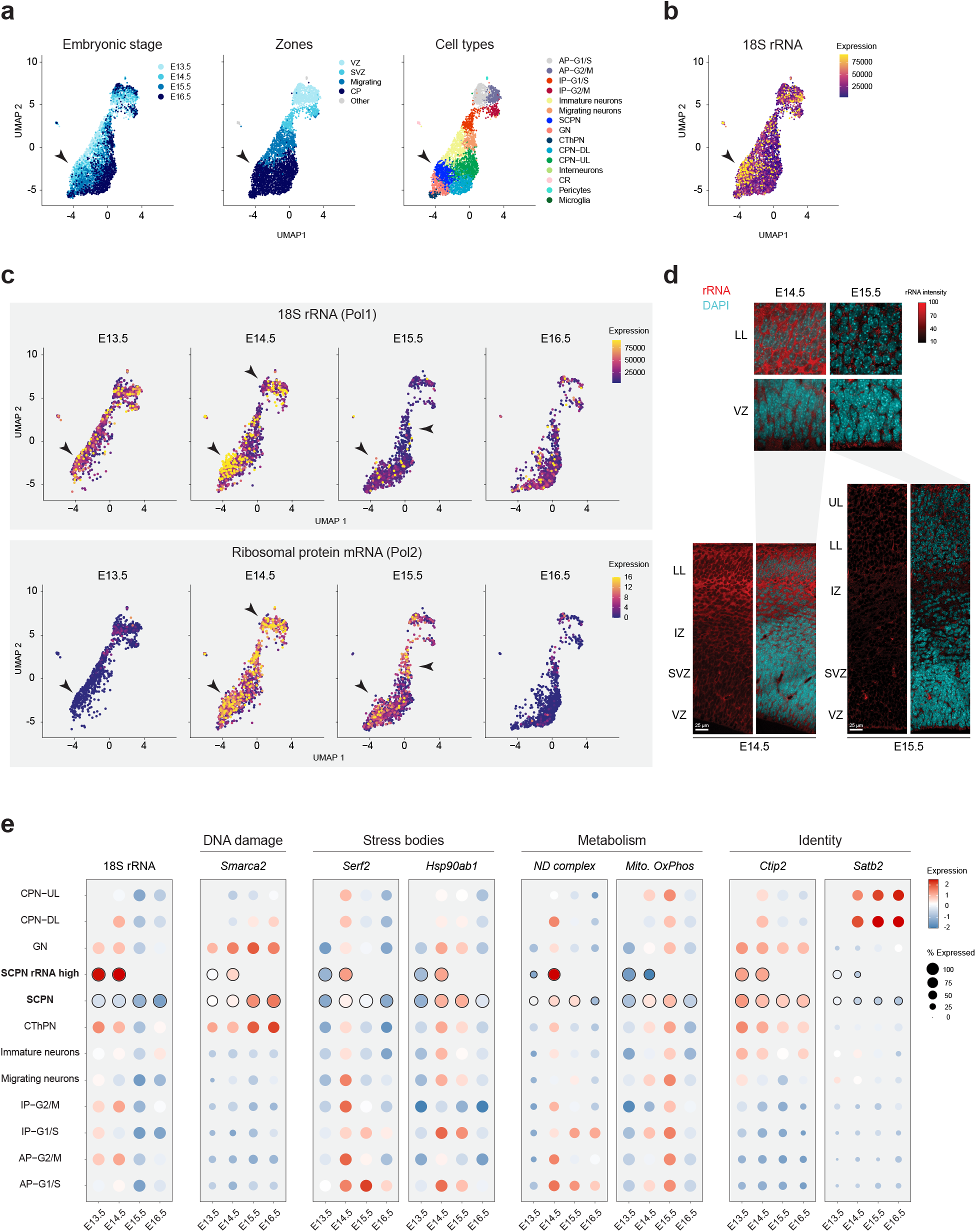
Neuronal subtype-specific rRNA transcription during cell state transitions. **a**: UMAP projection of 4186 mouse neocortex cells profiled with RiboSTS from E13.5 to E16.5 (zoomed in on 3890 cells; see **Extended Data Figure 10** for full map with interneurons and microglia). Cells are colored (from left to right) by embryonic stage, location in neocortex zones, and cell type annotation. Abbreviations: AP, apical progenitor; IP, intermediate progenitor; CThPN, corticothalamic projection neuron; GN, granular layer neuron; SCPN, subcerebral projection neuron; CPN, (intra)cortical projection neuron; CR, Cajal–Retzius cell; DL, deep layer; UL, upper layer, G1/S/G2/M, cell cycles phases. **b**: UMAP from (a) colored by 18S rRNA expression levels. Arrows indicate post-mitotic, post-migratory, SCPNs with high rRNA expression. **c**: UMAPs split by developmental stage and colored by 18S rRNA (top) or ribosomal protein mRNAs (bottom) expression. Ribosomal protein mRNA expression is the average of all core ribosomal protein mRNAs. **d**: Immunohistochemistry labeling of rRNA (red) and DAPI-stained DNA (cyan) in coronal neocortex sections at E14.5 and E15.5. Scale bar: 25 µm. VZ: ventricular zone; SVZ: subventricular zone; CP: cortical plate; IZ: intermediate zone; LL: lower layers; UL: upper layers. **e**: Expression of 18S rRNA in cell clusters over time, compared to genes in DNA damage, stress body, metabolism, and neuronal identity pathways.

The rRNA transcriptome demonstrates striking neuronal subtype and stage-specific production (**Fig. 4b**). Post-mitotic, post-migratory, lower layer subcortical projection neurons demonstrate particularly abundant rRNA production. These neurons are strongly *Ctip2* and *Fezf2* positive, *Tle4* and *Dcx* negative, characteristic of layer 5 subcerebral projection neurons (SCPN) that connect to the brainstem and spinal cord ^35,36^ (**Extended Data Fig. 10**). Elevated rRNA in SCPNs is transient, decreasing three-fold within 24 hours at E15.5 (**Fig. 4b-c**). These data indicate that rRNA transiently spikes during subcerebral projection neuron differentiation.

In contrast, later born, upper layer intracortical projection neurons demonstrate low rRNA production throughout their developmental trajectory (**Fig. 4b-c, Extended Data Fig. 10**). The rRNA decrease is already apparent in G1/S-phase apical progenitors at E14.5, particularly low in *Tbr2* positive intermediate progenitors that expand this lineage ^37^, and remain low in upper layer neurons destined for layers 4/3/2 expressing *Satb2* and *Cux1/2* ^38– 41^. Imaging the transition from E14.5-E15.5 by rRNA immunolabeling reinforces the sequencing data (**Fig. 4d**), with intense rRNA signal in lower layers at E14.5 followed by a sharp decrease at E15.5, and low levels throughout later born VZ progenitors and their daughter neurons. Low rRNA expression in intermediate progenitors expanding the upper layer lineage contrasts with high ribosomal protein mRNA expression in the same cells at E15.5 (**Fig. 4c**). These data highlight cell type-specific distinctions between Pol1 and Pol2 transcriptional programs, with rRNA production first increasing, then decreasing, ~24 hours ahead of ribosomal protein mRNAs.

Pathways associated with DNA damage, stress bodies, and metabolic change overlap the spike in rRNA production in SCPNs at E14.5 (**Fig. 4e**). DNA damage response factor *Smarca2* ^42,43^ is enriched lower layer neurons, and increases in SCPNS at E14.5. Stress body factors *Serf2* ^44^ and *Hsp90ab1* ^27^ are transiently elevated in nearly all cells at E14.5. Nuclear-encoded mitochondrial complex (ND) expression is strikingly elevated in rRNA-high SCPNs at E14.5, and uncoupled from the low expression of other oxidative phosphorylation (OxPhos) genes, indicative of a metabolic stress state ^45^. Taken together, an rRNA transcription spike occurs in SCPNs with distinct stress and metabolic profiles, which contrast with later upper layer lineages.

### Premature Pol1 shutdown reprograms neuronal identity

Finally, we tested the function of a transient peak in Pol1 activity for the fate of early-born lower layer lineages. shRNAs targeting *Polr1a* – the essential catalytic Pol1 subunit – were *in utero* electroporated into VZ progenitors at E12.5, and the fate of differentiating neurons analyzed at E16.5 (**Fig. 5a**). *Polr1a* knockdown had no observable effect on the migration or layer positioning of this lineage (**Fig. 5b**), in agreement with peak Pol1 activity occurring in post-mitotic, post-migratory SCPNs (**Fig. 4b-d**). However, immunostaining for subcerebral Ctip2 and intracortical Satb2 lineage markers revealed that *Polr1a* knockdown leads to a three-fold increase in the percentage of cells co-expressing both proteins (**Fig. 5c-d**). Ctip2-Satb2 double-positive cells typically represent 0-5% of differentiating mouse cortical neurons at E16.5, and acquire a distinct identity from SCPNs ^46,47^. Thus, timed nucleolar Pol1 activity directs neuronal subtype specification in the neocortex (**Fig. 5e**).

**Figure 5.**
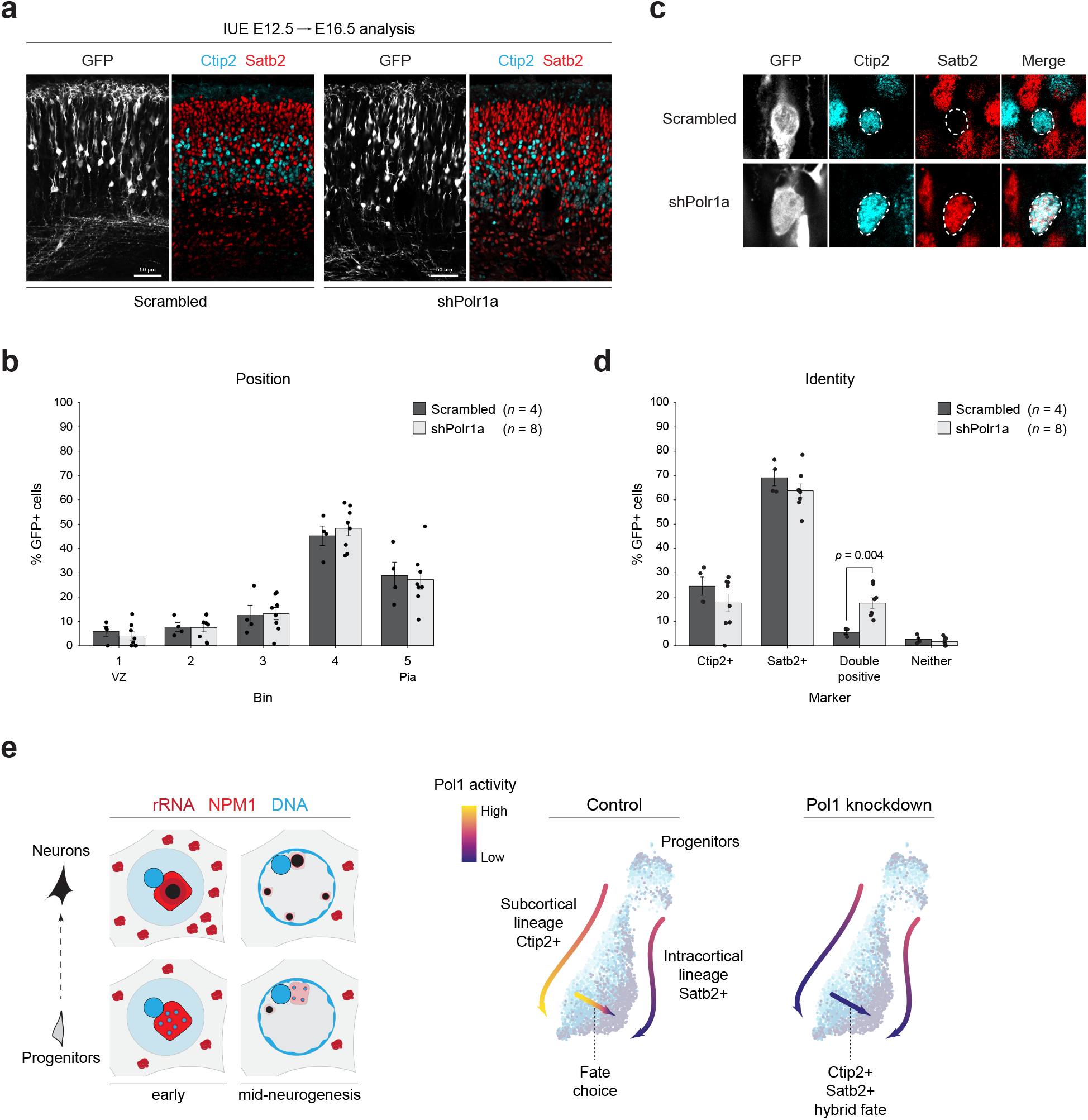
Pol1 knockdown in the rRNA abundant lower layer lineage redirects cell fate. **a**: Electroporation of shRNAs targeting *Polr1a* or scrambled control *in utero* at E12.5, with histology and microscopy analysis shown at E16.5. All electroporated cells are labeled with GFP. Immunostaining for lower layer marker Ctip2, and upper layer marker Satb2. **b**: Quantification of electroporated cell position from (a) in five bins spanning the ventricular zone progenitor niche (VZ, Bin 1), to the surface above differentiating neurons in the cortical plate (Pia, Bin 5). **c**: Images of Ctip2 and Satb2 immunostaining in electroporated cells, illustrating exclusive or co-expression. **d**: Quantification of (c) for percent of electroporated cells expressing Ctip2, Satb2, both (double positive), or neither. **e**: Summary of nucleolar architecture, rRNA transcription, and Pol1 function during neuronal subtype specification in neocortex development.

## Discussion

Our findings build upon prior discoveries of transcription and translation regulation in the neocortex ^1^. Developmental cues stimulate transcription factor binding to guide Pol2 activity, generating the mRNA transcriptomes of neocortex progenitors and neurons ^2,3,5,6^. Simultaneously, layers of post-transcriptional regulation, including translation by the ribosome, pattern neuronal subtype-specific proteomes ^1,48–50^. Recent work highlights how multi-level transcriptome and proteome expression vary in neocortex development ^4,51^, but how mRNA transcription and translation are coordinated remains unclear. The nucleolus is a hub that integrates information from developmental cues and metabolic states to guide Pol1 activity, rRNA transcription, and ribosome biogenesis – generating the mRNA translation capacity of each cell ^8–10^. We uncover dynamic nucleolar architectures and dual rRNA-mRNA transcriptome states that define neuronal subtypes in the neocortex.

Our study advances several important concepts in the molecular model of neurodevelopment (**Fig. 5e**). The nucleolus is not a consistent, housekeeping organelle during neurogenesis; but rather, it displays unique features in progenitors and neuronal subtypes over developmental time. Neurons accumulating in lower layers at E14.5 have nucleoli restructured with a single central cavity, which is associated with rDNA transcription stress more commonly seen in non-mammalian species ^9,21–28^. Nucleolar cavities can harbor rRNA surveillance complexes like the exosome, and exclude ribosomal proteins and assembly factors ^21,52^. Hypertranscription of rDNA in lower layer lineages may trigger the formation of nucleolar cavities as Npm1 is depleted, and lead to nucleolus silencing at the nuclear periphery ^19^. The mobilization of nucleolus-associated chromatin to the periphery may impact the activity large genomic regions during neurogenesis ^53–55^. Nucleolar cavities can also sequester Pol2 and spliceosome proteins ^21,56,57^, raising the intriguing possibility that the nucleolus can directly engage crosstalk between Pol1 and Pol2 ^14,58^ during neurogenesis.

Simultaneous analysis of rRNA and mRNA transcriptomes with RiboSTS links nucleolar dynamics to neuronal subtypes and cell states (**Fig. 5e**). rRNA production spikes in specific neurons after migration at E14.5, which are destined to form corticospinal projections and express Ctip2 ^36^. Shutdown of Pol1 activity in this lineage generates a hybrid identity at E16.5, co-expressing Ctip2 with Satb2 ^38,39^, which is typically expressed in the subsequent rRNA-depleted lineage. Double-positive Ctip2-Satb2 neurons are normally scarce at E16.5 and increase later, forming projections to either the subcortical brainstem or contralateral cortex ^46,47^. Despite the ubiquitous requirement for Pol1, its activity is heterogeneous, and Pol1 mutations lead to tissue-specific phenotypes that include neurodevelopmental abnormalities in humans ^30,59–61^.

In summary, transcription and translation regulation converging in the nucleolus may be a key feature of neurodevelopment. We anticipate that RiboSTS will spark the discovery of dual rRNA and mRNA transcriptome heterogeneity in diverse biological systems and disease states at single-cell resolution.

## Competing interests

RiboSTS has been filed under patent number EP25184315.7 (pending): Single Cell Analysis of Structural RNA (2025).

## Figure Legends

**Extended Data Figure 1.**
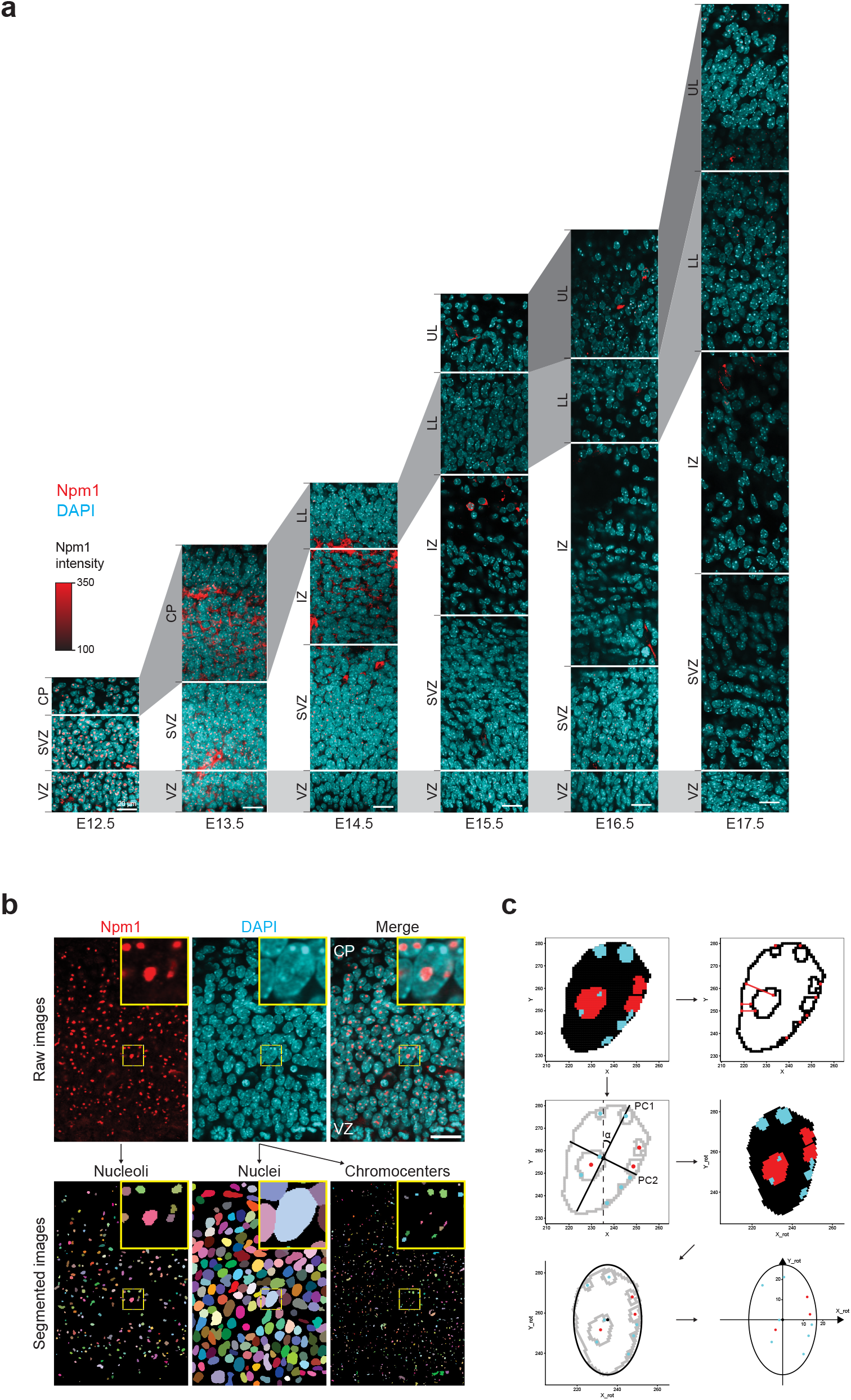
Image quantification workflow. **a**: Immunohistochemistry labeling of Npm1 (red) and DAPI-stained DNA (cyan) in coronal neocortex sections with layer annotations. Scale bar: 20 µm. Abbreviations: VZ: ventricular zone; SVZ: subventricular zone; CP: cortical plate; IZ: intermediate zone; LL: lower layers; UL: upper layers. **b**: Representative images showing segmentation output of the Npm1 channel (nucleolus mask), and DAPI channel (nucleus and chromocenter masks). **c**: Schematic of the workflow for extracting distance information. Edge-to-edge distances (shown as red lines) were calculated for nucleolus-nucleus, chromocenter-nucleus, and nucleolus-chromocenter. Nucleoli are shown in red and chromocenters in blue. For plotting center-to-center distances, the longest axis of each nucleus (PC1 axis) was aligned to the vertical axis of the image by rotating it through the smallest angle (α). An elliptical approximation each nucleus’s area was constructed using the PC1 and PC2 axes.

**Extended Data Figure 2.**
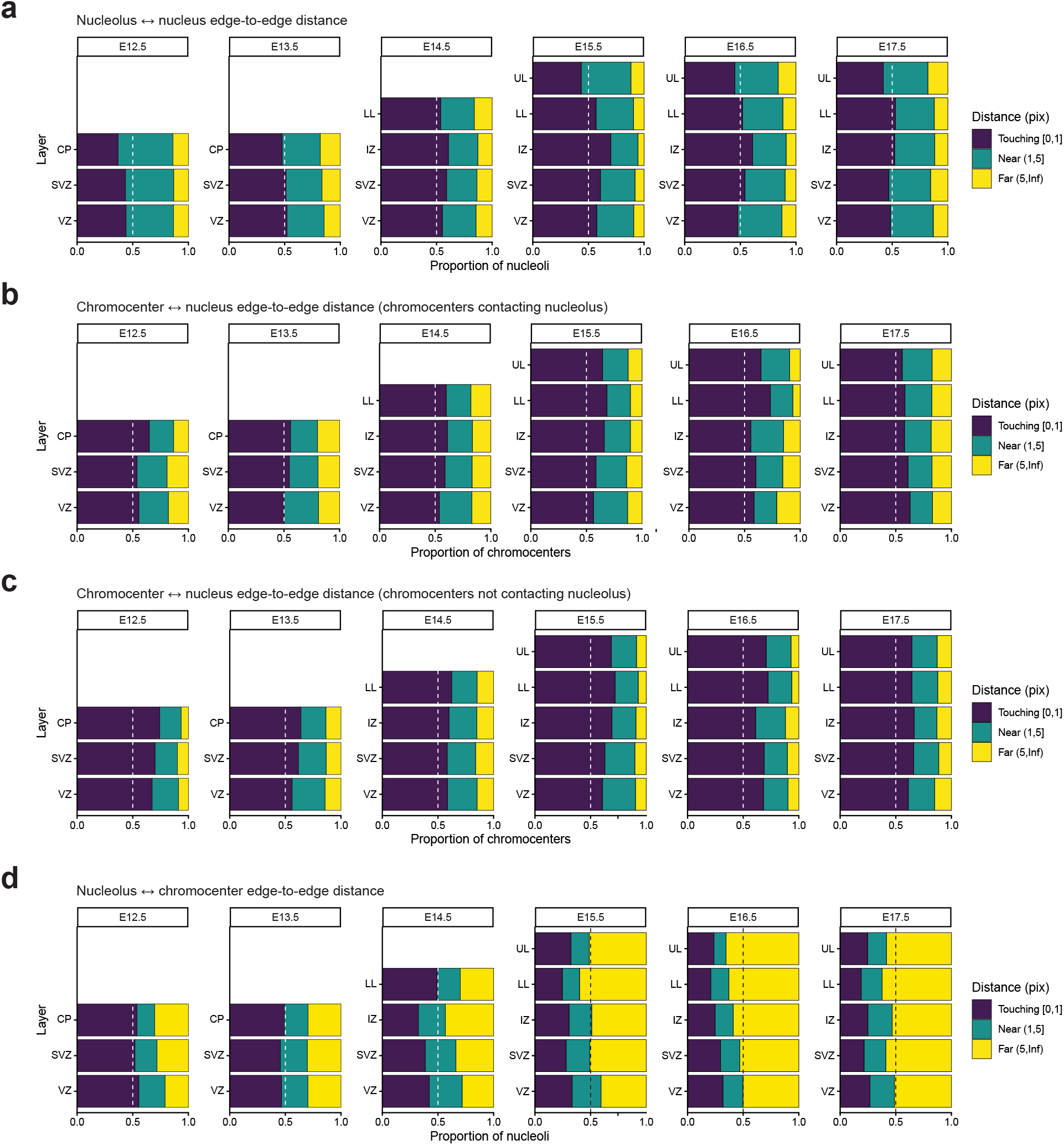
Nucleolus-chromocenter contacts in the neocortex across developmental stages. Stacked barplot quantification of edge-to-edge distances by age and layer, categorized into three distance categories: “Touching” (0–1 pixel or 0–0.22 µm), “Near” (1–5 pixels or 0.22– 1.10 µm), or “Far” (>5 pixels or >1.10 µm): **a**: Nucleolus-to-nucleus distances. **b**: Chromocenter-to-nucleus distances for chromocenters contacting nucleoli. **c**: Chromocenter-to-nucleus distances for chromocenters not contacting nucleoli. **d**: Nucleolus-to-chromocenter distances.

**Extended Data Figure 3.**
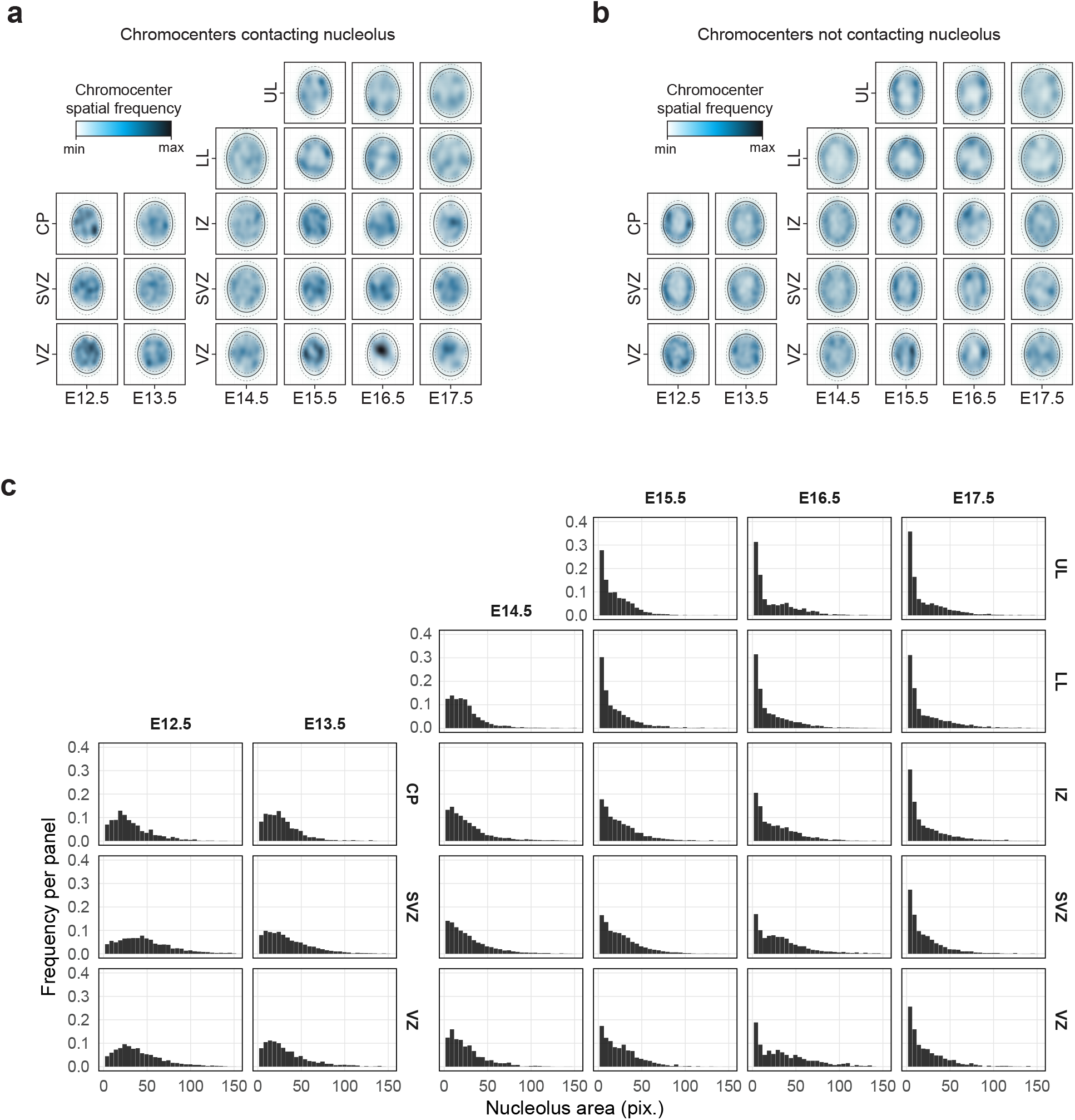
Spatial distribution of chromocenters with and without nucleolus contacts. **a**: Spatial distribution of nucleoli-contacting chromocenters within the nucleus relative to the nuclear center of mass, with elliptical approximation of nuclear area. The solid line represents the median, and dashed lines indicate the 25th and 75th percentiles. Kernel density estimates are depicted (white: low density; dark blue: high density), normalized within each panel. **b**: Similar spatial distribution quantification for nucleoli-non-contacting chromocenters. **c**: Histogram quantification of nucleolar area, represented as pixel number (pix.), across stages and layers.

**Extended Data Figure 4.**
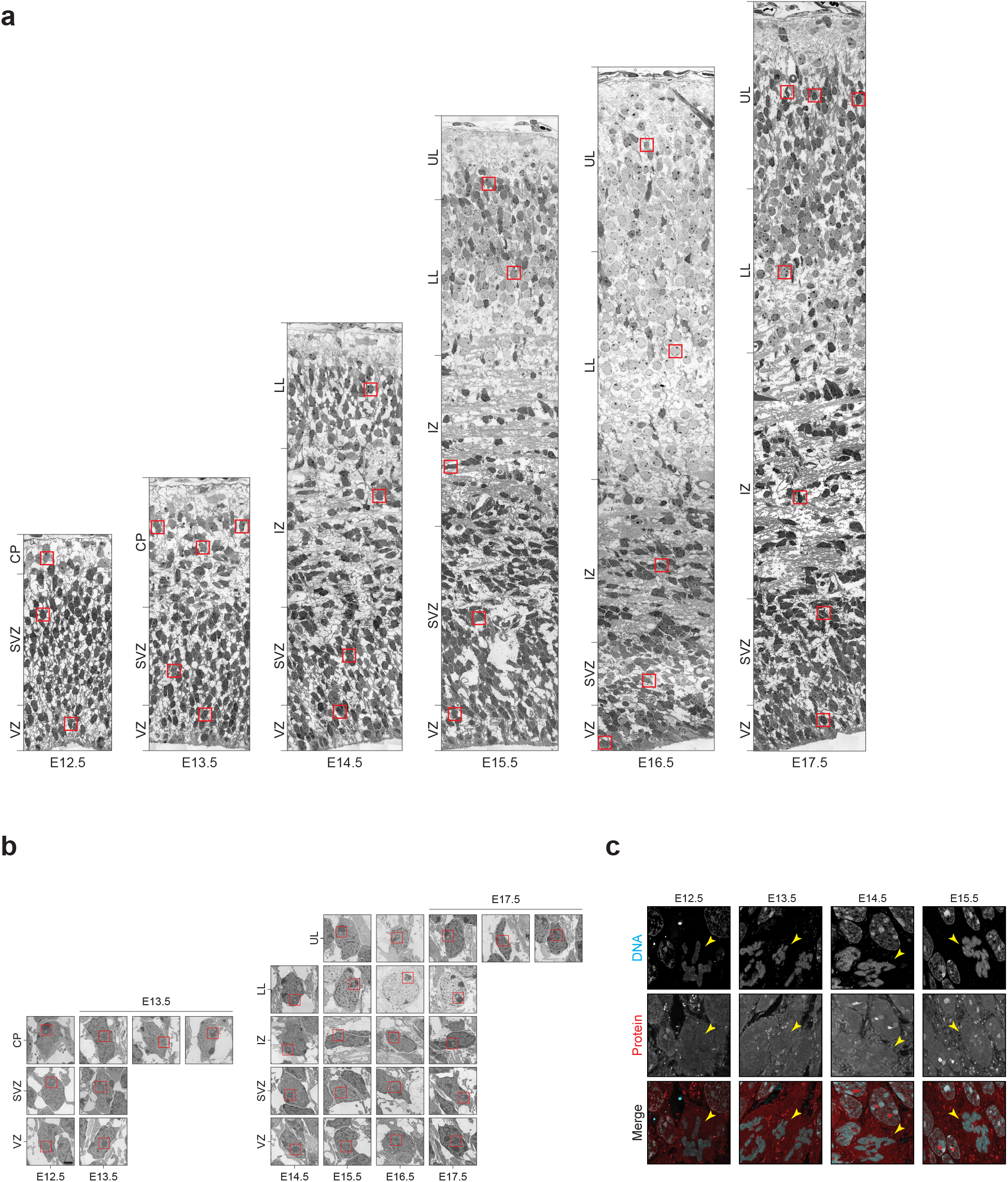
Additional data for electron and expansion microscopy. **a**: Overview of coronal neocortex sections imaged with electron microscopy, with layer annotations. Red squares indicate the cells that are show in Figure 2a. Abbreviations: VZ: ventricular zone; SVZ: subventricular zone; CP: cortical plate; IZ: intermediate zone; LL: lower layers; UL: upper layers. **b**: Zoomed-in view of the nuclei for cells annotated with a red square in (a). Scale bar: 2 µm. **c**: Mitotic cells lacking nucleoli (serving as a negative control for nucleolus identification) visualized with expansion microscopy in coronal neocortex sections with labeling of total protein (red) and DNA (cyan), associated with Figure 2b. Yellow arrows highlight condensed chromosomes.

**Extended Data Figure 5.**
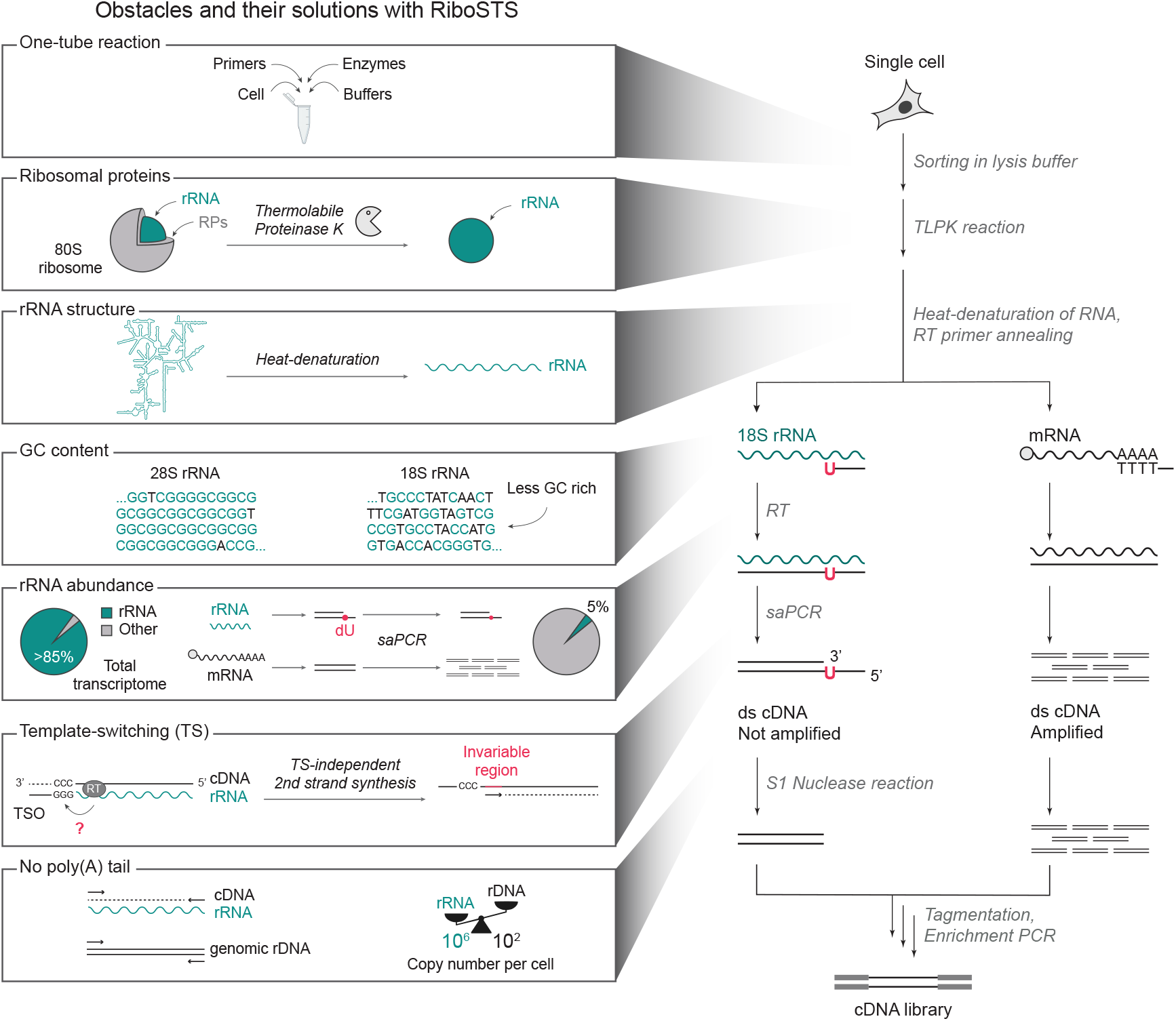
Obstacles and their solutions with RiboSTS. Flowchart of the RiboSTS method, including the obstacles addressed by RiboSTS at each step (left) and the key innovations introduced to overcome them (right).

**Extended Data Figure 6.**
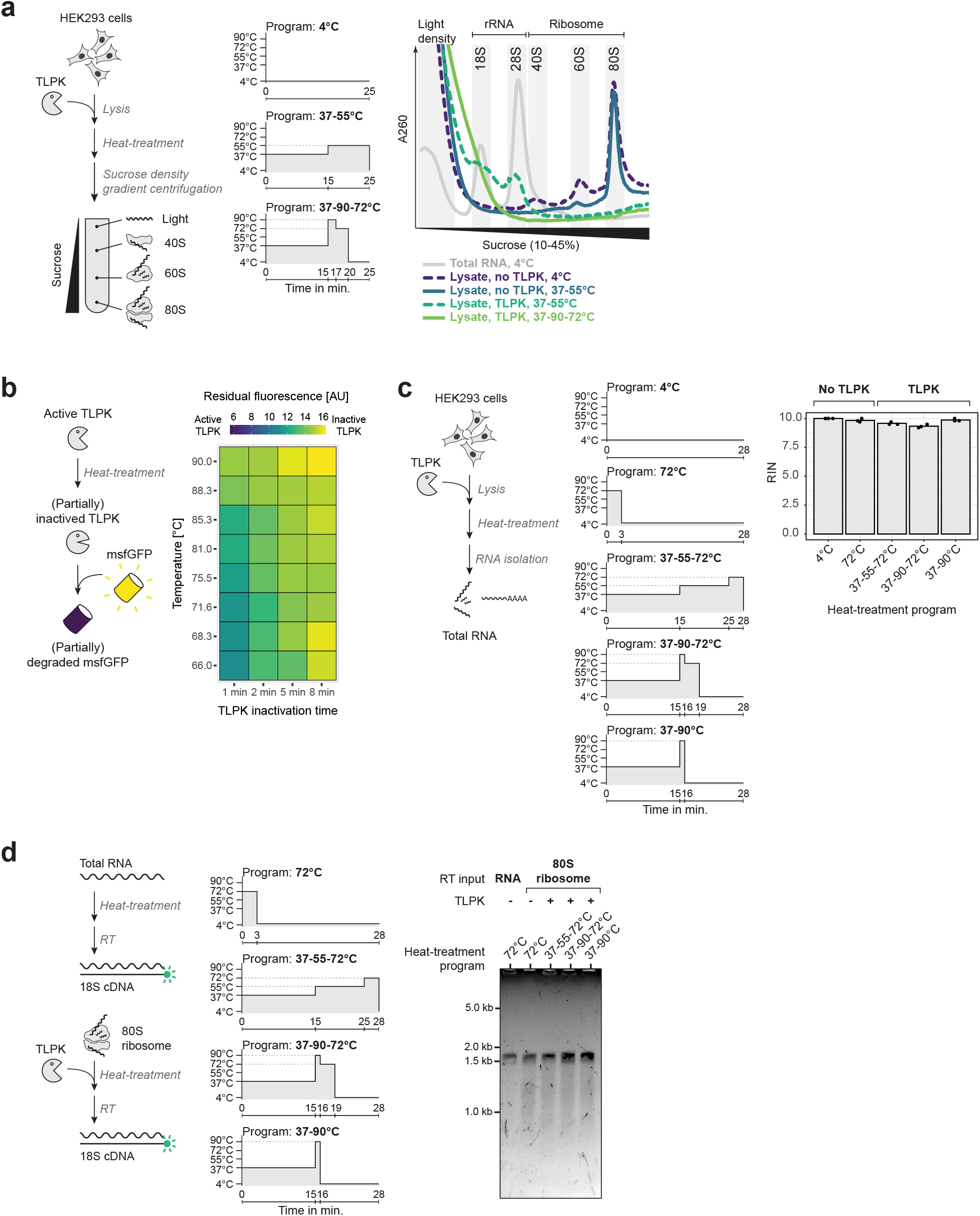
Optimization of TLPK treatment for compatibility with a one-tube single-cell sequencing reaction. **a**: Thermolabile Proteinase K (TLPK) degrades ribosomal complexes and releases rRNA. Experimental schematic with heat-treatment procedures (left), followed by sucrose density gradient analysis (right). **b**: Optimization of the TLPK heat-inactivation protocol. Active TLPK was heat-treated, then incubated with msfGFP to assess residual activity via fluorescence. **c**: RNA integrity after TLPK treatment and heat-inactivation. Right: RNA integrity number (RIN) analysis (n = 3). **d**: Reverse transcription (RT) of 18S rRNA after TLPK treatment, with cDNA yield analyzed by alkaline agarose gel electrophoresis.

**Extended Data Figure 7.**
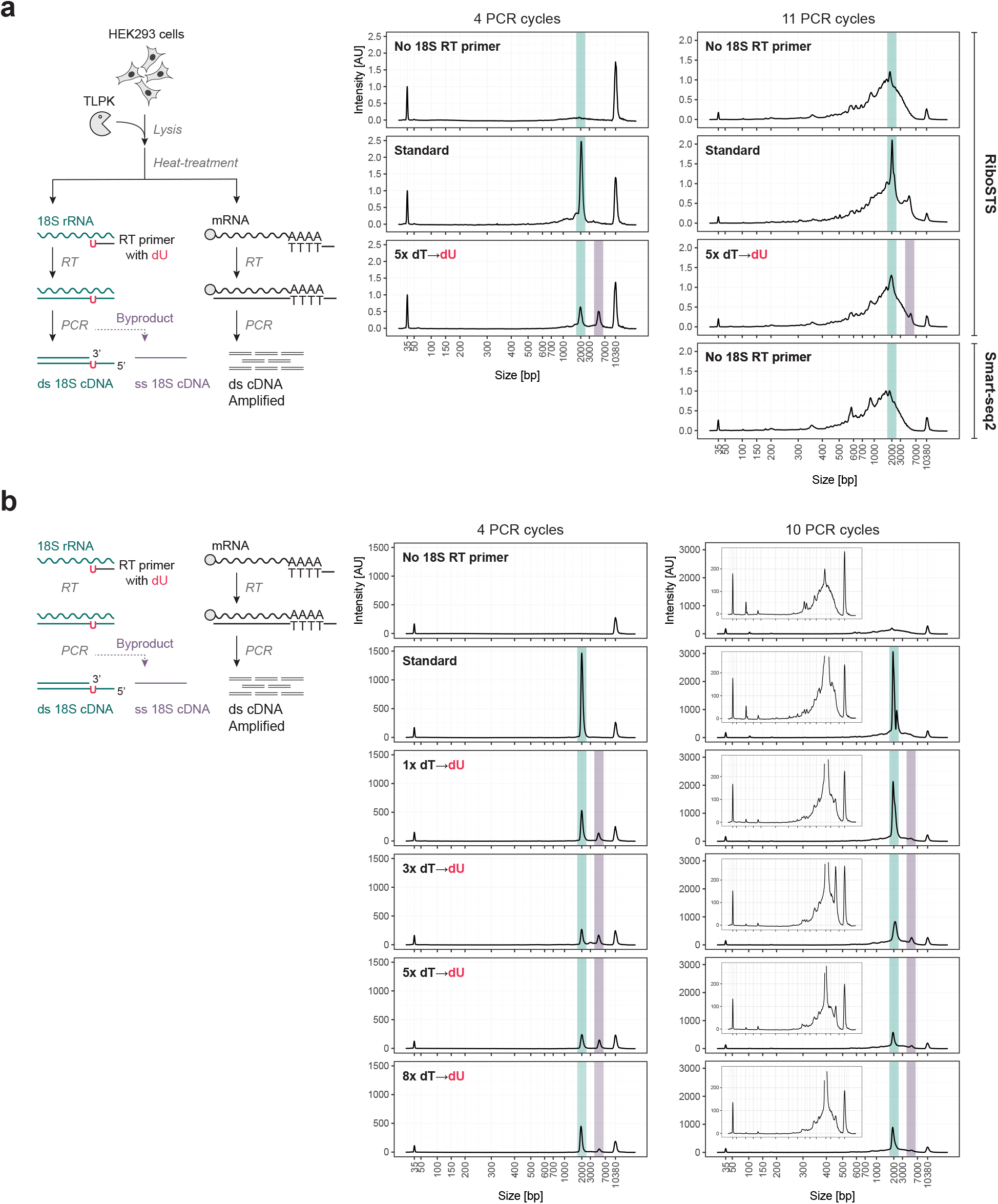
Balancing cDNA yield from 18S rRNA and mRNA through selective amplification PCR (saPCR). **a**: cDNA balancing initiated from cell lysates with varying numbers of dT-to-dU substitutions in the RT primer, with increasing PCR cycles. Experimental schematic (left), followed by analysis of saPCR products on Bioanalyzer (right). Double-stranded 18S cDNA is shown in green, byproduct single-stranded 18S cDNA is shown in purple. A Smart-seq2 sample, prepared without TLPK, using only an RT primer targeting mRNA, served as a comparison. **b**: cDNA balancing initiated from purifying total RNA omitting the TLPK treatment, with varying dT-to-dU substitutions in the RT primer across increasing PCR cycles.

**Extended Data Figure 8.**
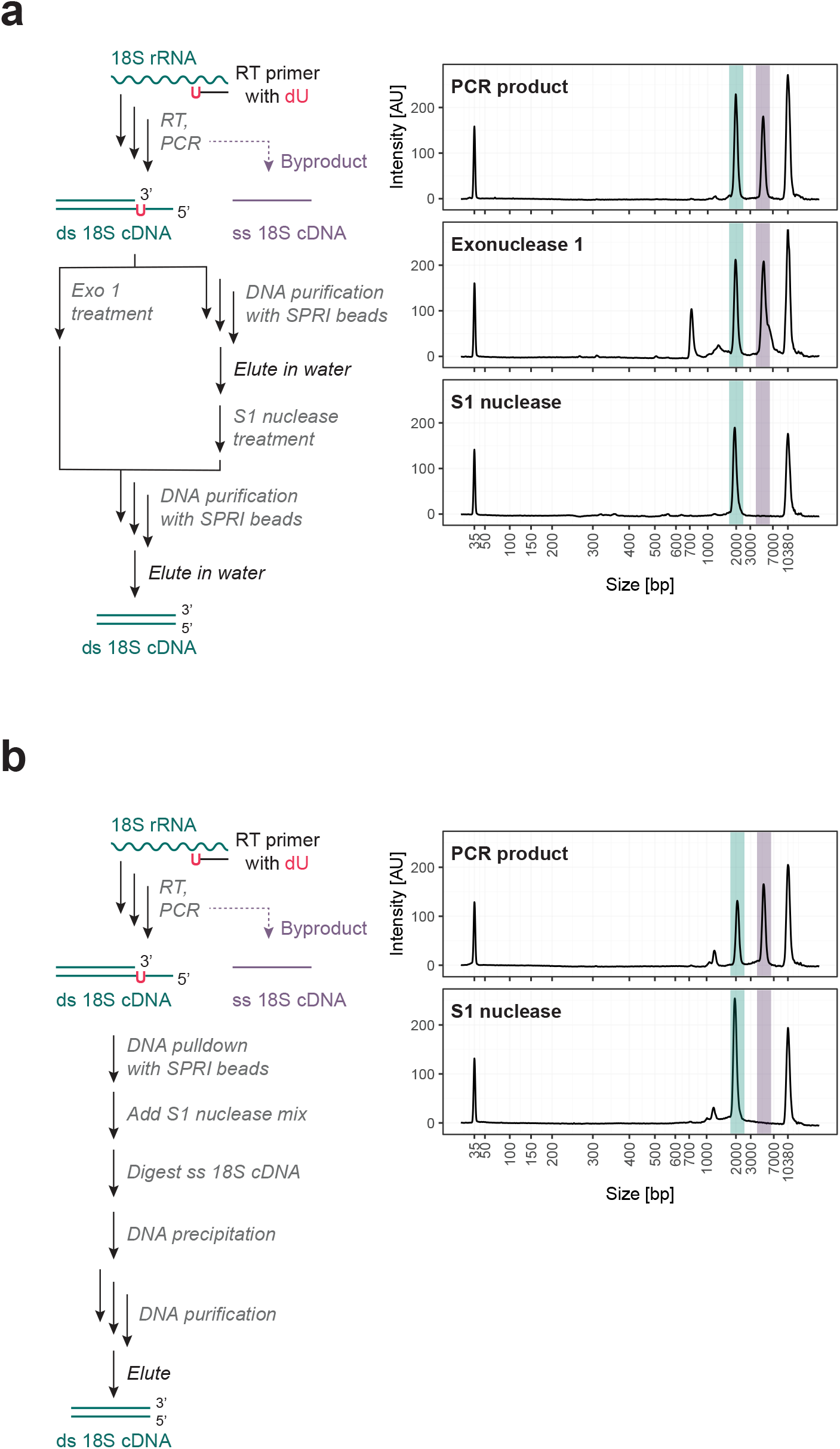
Removal of saPCR byproducts in a one-tube reaction. **a**: Comparison of nuclease treatments on the removal of ss 18S cDNA saPCR byproducts. S1 nuclease selectively removes the saPCR balancing byproduct, unlike Exonuclease 1. Experimental schematic (left), followed by analysis of saPCR products after nuclease treatment on Bioanalyzer (right). **b**: Removal of single-stranded 18S cDNA (ss18S) byproduct using DNA pulldown with SPRI beads, followed by resuspension in an S1 nuclease reaction mix. This method enables the simplification to a single, scalable purification step for eliminating the ss18S cDNA byproduct (indicated by the purple peak). Experimental schematic (left), followed by analysis of saPCR products after S1 nuclease treatment on Bioanalyzer (right).

**Extended Data Figure 9.**
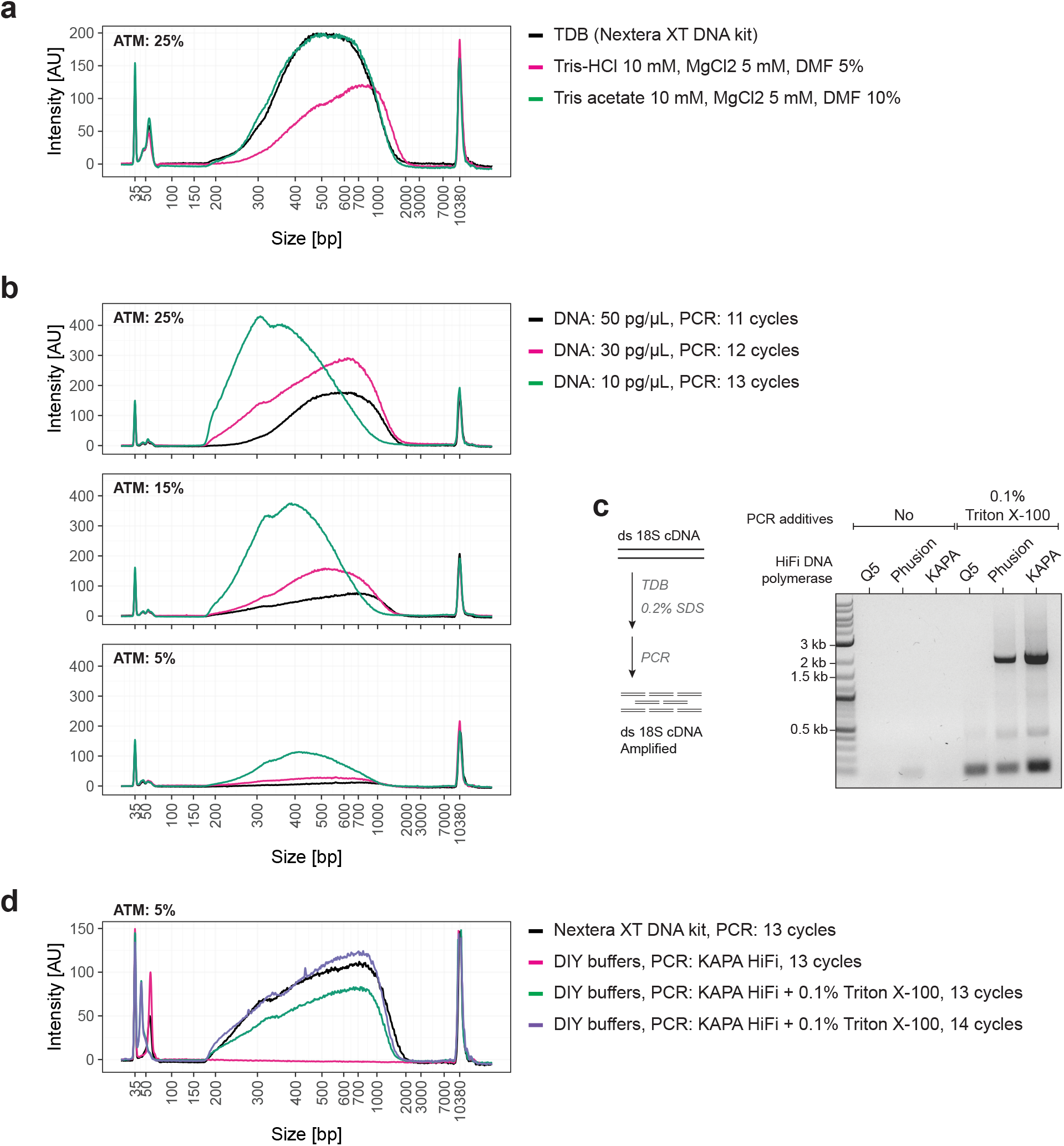
A more cost-efficient tagmentation reaction with custom buffers. **a**: Determining the composition of a substitute tagmentation reaction buffer for TDB from the Nextera XT DNA Kit. **b**: Reducing the concentration of ATM requires a proportional decrease in DNA concentration for effective tagmentation. **c**: The inhibitory effect of SDS on KAPA HiFi DNA polymerase can be countered by adding Triton X-100 to the PCR mix. **d**: The efficiency of a miniaturized tagmentation reaction using custom buffers matches that of commercial reagents.

**Extended Data Figure 10.**
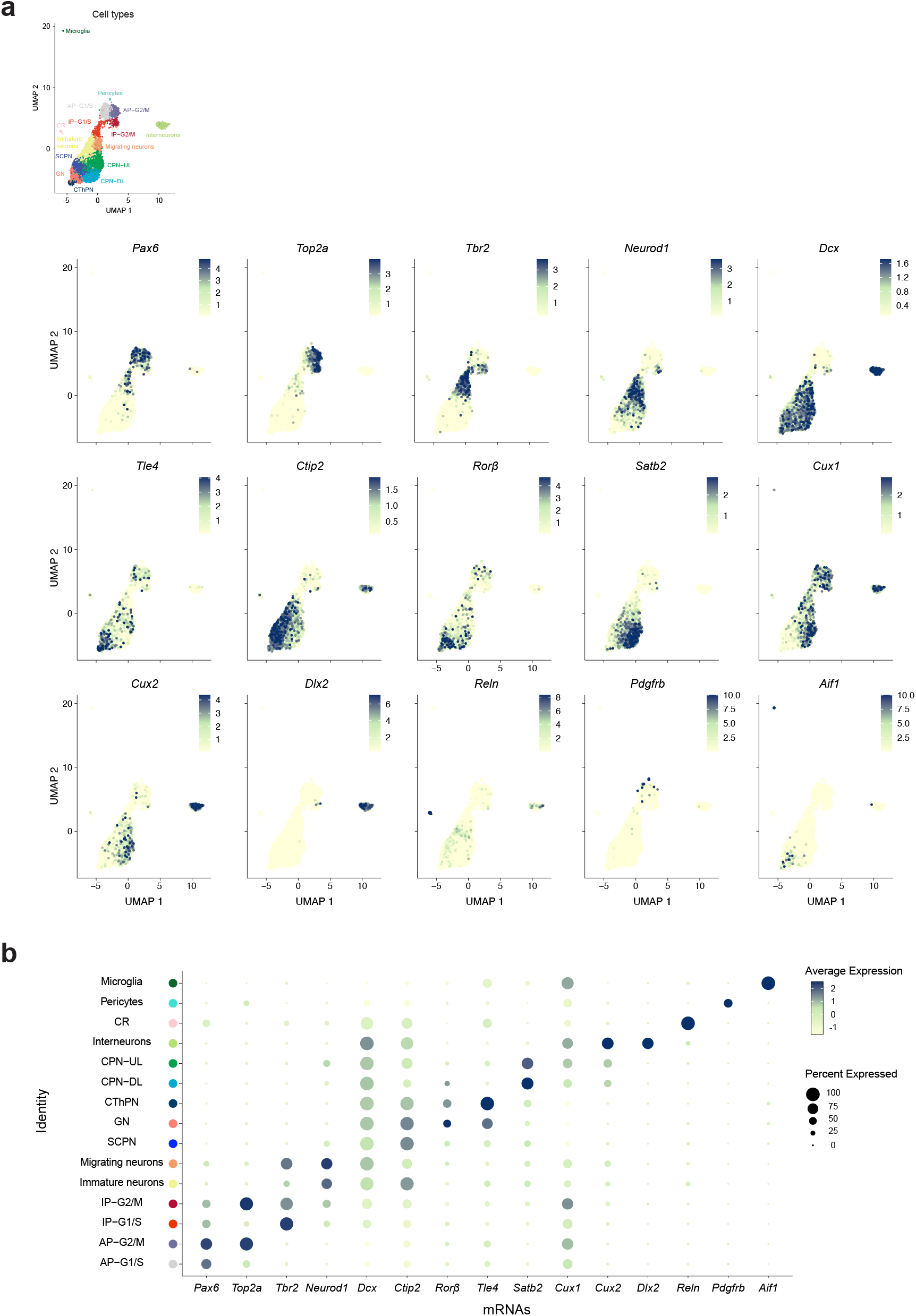
Expression of marker genes and neocortex cell type annotation. **a**: Combined UMAP of E13.5 to E16.5 cells colored by the expression of representative marker genes defining cell clusters. Apical progenitors (*Pax6*) G2/M phase (*Top2a*), basal intermediate progenitors (*Tbr2*), differentiating neurons (*Neurod1*), migrating neurons (*Dcx*), corticothalamic layer 6 projection neurons (*Tle4*), corticospinal layer 5/6 projection neurons (*Ctip2*), granular layer neurons (*Rorβ*), intracortically projecting upper layer 2/3/4 neurons (*Satb2, Cux1, Cux2*), interneurons (*Dlx2*), Cajal-Retzius cells (*Reln*), microglia (*Aif1*), and pericytes (*Pdgfrb*). **b**: Dot plot for the expression of marker genes in (a) in the cell clusters annotated in Figure 4. Abbreviations: AP, apical progenitor; IP, intermediate progenitor; CThPN, corticothalamic projection neuron; SCPN, subcerebral projection neuron; GN, granular layer neuron; CPN, (intra)cortical projection neuron; CR, Cajal-Retzius cell; DL, deep layer; UL, upper layer, G1/S/G2/M, cell cycles phases.

## Notes

### Funding

- Deutsche Forschungsgemeinschaft: KR 5755/3-1, KR 5755/1-1

